# *Saccharomyces boulardii* attenuates obesity-associated inflammation and weight gain through coordinated gut ecosystem remodeling

**DOI:** 10.64898/2026.04.01.715546

**Authors:** Karl Alex Hedin, Troels Holger Vaaben, Ditte Olsen Lützhøft, Benjamin A. H. Jensen, Morten Otto Alexander Sommer

## Abstract

The gut microbiome is a key regulator of metabolic homeostasis and contributes to obesity progression through effects on immune signaling, gut barrier integrity, and systemic inflammation. Microbiome-targeted strategies are therefore being explored as complementary approaches to conventional weight-loss therapies. Here, we investigated the probiotic yeast *Saccharomyces boulardii* in a murine model of diet-induced obesity (DIO) using an integrated multi-omics framework combining metabolic phenotyping, gut microbiome profiling, cecal metabolomics, colonic transcriptomics, and portal cytokine analysis. *S. boulardii* reduced food intake, attenuated weight gain, and increased energy expenditure without major changes in circulating metabolic hormone levels. Microbial diversity remained largely preserved, but selective enrichment of Bacteroidales lineages, including Muribaculaceae, was observed alongside functional remodeling of microbial pathways. Cecal metabolomics revealed increased B-vitamins, betaine, and GABA, with reduced stress-associated metabolites. Colonic transcriptomics showed attenuation of TNFα/NF-κB signaling and enrichment of interferon and epithelial programs, while portal cytokine profiling indicated reduced inflammatory chemokines with trends toward increased IL-17A and IL-22. Integrated multi-omics analysis identified coordinated host–microbe interactions across metabolic, transcriptional, and immune layers. Collectively, these findings demonstrate that *S. boulardii* modulates the gut–immune–metabolic axis in obesity, supporting microbiome-based interventions as potential adjunct strategies targeting metabolic inflammation.

## Introduction

The gut microbiome is a central regulator of metabolic homeostasis and a key driver of obesity progression through its effects on host immunity, gut barrier integrity, and systemic inflammation^1^. Accumulating evidence suggests that commensal microbes influence inflammatory tone, energy balance, glucose handling, and lipid metabolism, supporting a role for the gut microbiome as an upstream contributor to metabolic dysfunction^2^. Microbiome configurations associated with metabolic health are characterized by enhanced gut barrier function and reduced inflammatory signaling^3^. Consistent with this concept, fecal microbiota transfer (FMT) from lean donors attenuates weight gain and improves metabolic parameters in preclinical obesity models^4^. However, translation to humans has yielded more variable outcomes, with some studies reporting improvements in insulin sensitivity but limited or no effects on body weight or long-term metabolic parameters^5,6^. These findings highlight the complexity of host–microbiome interactions in humans and the need for more targeted microbiome-based interventions. Accordingly, microbiome-targeted modulation strategies are increasingly being explored in clinical settings^7^.

Importantly, experimental manipulation of the gut microbiota can markedly influence host metabolism. In some diet-induced obesity (DIO) models, depletion of the microbiota through antibiotic treatment or germ-free conditions improves insulin sensitivity, enhances GLP-1 secretion, and reduces hyperinsulinemia^8^. These findings highlight the microbiota as an active regulator of host energy metabolism. However, the relationship between microbiota composition and metabolic health is context-dependent, as a balanced microbial community is also critical for maintaining metabolic homeostasis and immune regulation^9,10^. Collectively, these insights have driven the development of microbiome-targeted interventions for obesity, including probiotics, prebiotics, FMT, and advanced microbiome therapeutics (AMTs)^11–13^, with the aim of restoring immune–metabolic homeostasis rather than solely suppressing appetite or caloric intake.

Among these approaches, selected probiotic strains have shown promise in modulating host immunity, gut barrier integrity, and microbiota composition in obesity^11,14^. For example, supplementation with *Bifidobacterium breve* B-3 reduced body fat in pre-obese adults^15^, while pasteurized *Akkermansia muciniphila* improved insulin sensitivity with only modest effects on body weight^16^, highlighting dissociation between metabolic benefits and weight loss. The probiotic yeast *Saccharomyces boulardii* has been reported to reduce body weight and fat mass in db/db mice^17^. In obese adults (BMI 30–35 kg/m²), a randomized, double-blind, placebo-controlled trial evaluating a 60-day supplement containing *S. boulardii* and superoxide dismutase (SOD) demonstrated reductions in body weight, BMI, and fat mass with preservation of lean mass^18^; however, the inclusion of SOD complicates attribution of these effects specifically to *S. boulardii*.

Beyond effects on adiposity, *S. boulardii* exhibits biological properties directly relevant to obesity-associated inflammation. It exerts antimicrobial activity, modulates innate and adaptive immune responses, and enhances the production of short-chain fatty acids (SCFAs), which are critical for intestinal barrier integrity and immune regulation^19^. Notably, *S. boulardii* strengthens epithelial barrier function^20^ and reduces circulating lipopolysaccharide (LPS) levels—a key driver of metabolic endotoxemia and chronic low-grade inflammation in obesity^21^. Despite these pleiotropic effects, the mechanisms by which *S. boulardii* influences host metabolism, particularly through inflammation-dependent pathways, remain insufficiently defined.

In this study, we investigated the effects of *S. boulardii* in a murine model of established DIO with a specific focus on inflammation-linked metabolic dysfunction. We employed an integrated multi-omics approach combining gut microbiome profiling, metabolic phenotyping, colonic transcriptomics (RNA-seq), and systemic cytokine measurements to examine host–microbe–immune interactions. This design allowed us to delineate how *S. boulardii*-driven alterations in the gut ecosystem translate into changes in intestinal inflammatory signaling and downstream systemic immune tone, independent of primary appetite suppression or weight loss. By coupling microbial, tissue-level, and circulating immune readouts, our study provides mechanistic insight into how microbiome-based interventions may ameliorate obesity-associated inflammation and metabolic dysregulation as an adjunct to existing therapeutic strategies.

## Results

### Oral administration of *S. boulardii* attenuates weight gain and alters energy balance in DIO mice

Male C57BL/6 mice were maintained on a high-fat diet (HFD) for 8 weeks to establish DIO before receiving daily oral gavage of phosphate-buffered saline (PBS) or *S. boulardii* for an additional 7 weeks (Figure 2A). *S. boulardii* treatment significantly reduced cumulative food intake over the 40-day intervention period (Figure 2B), corresponding to an approximately 10% decrease in mean daily food consumption (Figure 2C).

**Figure 1.**
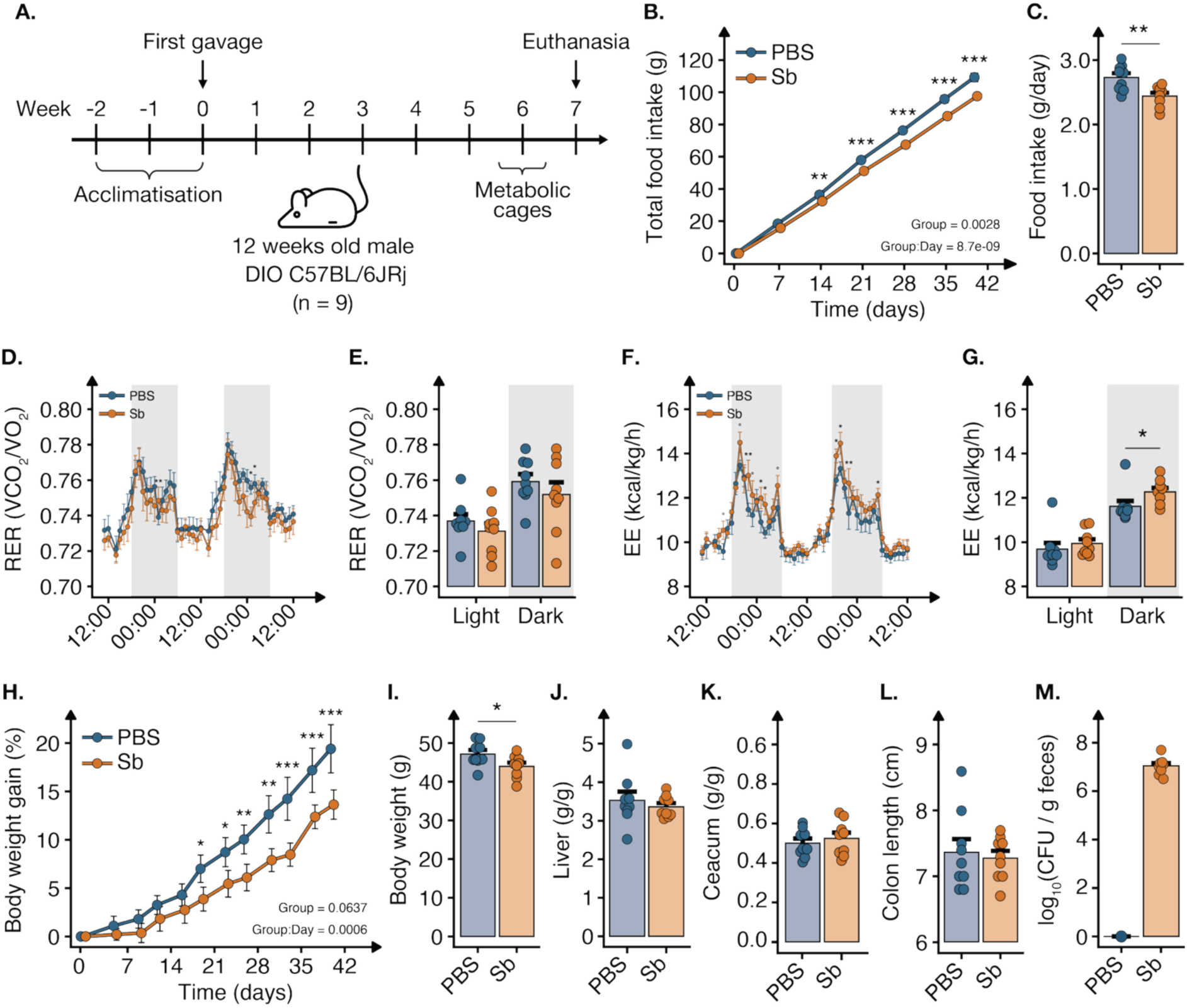
*S. boulardii* alters energy balance and attenuates weight gain in mice with DIO. **(A)** Study design; Male C57BL/6 mice with DIO 8 weeks) received oral PBS or *S. boulardii* while maintained on a high-fat diet (HFD; 60% kcal fat) for 7 weeks. **(B)** Cumulative food intake (g) over 40 days, measured weekly. **(C)** Mean daily food intake (g/day). **(D)** Hourly respiratory exchange ratio (RER; VCO_2_/VO_2_) over a 2-day period under a 12:12 light–dark cycle. **(E)** Mean RER during light and dark phases. **(F)** Hourly energy expenditure (EE; kcal/kg/h) over a 2-day period under a 12:12 light–dark cycle. **(G)** Mean EE during light and dark phases. (H) Body weight gain (%) over 40 days, measured twice weekly. (I) Body weight at endpoint (g). (J) Lever weight (g/g) and (K) total cecal weight (g/g) normalized to body weight. (L) Colon length (cm) at endpoint. (M) Log_10_ colony forming unities of *S. boulardii* per gram feces. Data are mean ± SEM (n = 9). Statistical analyses were performed using a linear mixed-effects model fitted with lmer (REML = FALSE), including Group, Day, and their interaction as fixed effects and mouse as a random intercept. Fixed effects were evaluated using Type III ANOVA (panels B, H); a two-tailed Student’s t-test (panels C, E, G, I–L); \**p* < 0.05, \*\**p* < 0.01, *** *p* < 0.001.

**Figure 2.**
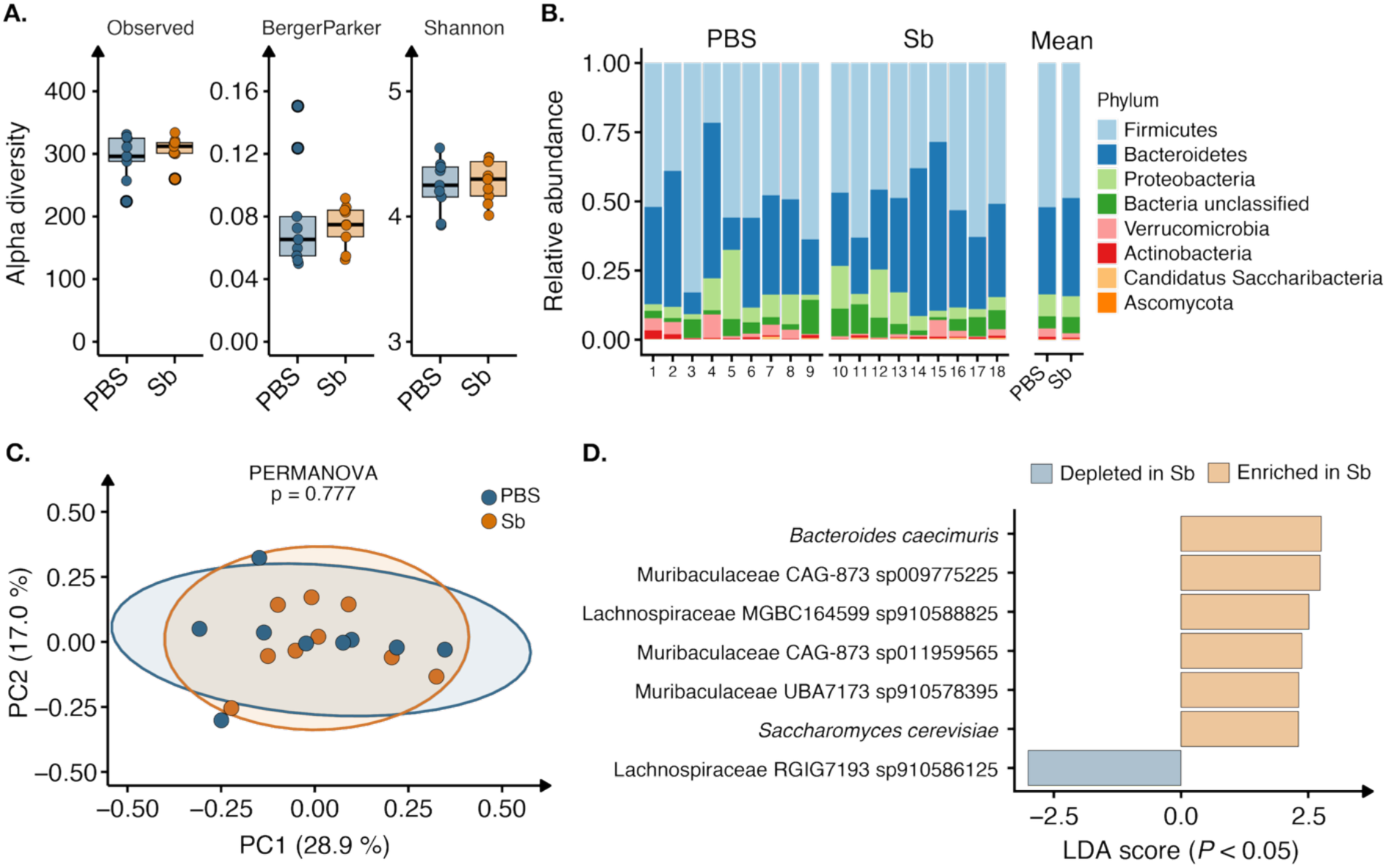
Gut microbiome composition following *S. boulardii* treatment in diet-induced obese mice (DIO). **(A)** Alpha diversity metrics including observed richness, Berger–Parker dominance, and Shannon diversity index in PBS- and *S. boulardii*–treated mice. **(B)** Relative abundance of microbial phyla in PBS- and *S. boulardii*–treated mice. **(C)** Principal coordinate analysis (PCoA) of beta diversity based on Bray–Curtis dissimilarity. Each point represents an individual mouse. Statistical significance was assessed using PERMANOVA. **(D)** Differentially abundant microbial species identified using linear discriminant analysis effect size (LEfSe), displayed as LDA scores indicating enrichment in PBS- or *S. boulardii*–treated groups. Group differences were assessed using two-sided Kruskal–Wallis and Wilcoxon rank-sum tests with Benjamini–Hochberg false discovery rate (FDR) correction (*p* < 0.05). Taxa with LDA scores > 2.0 were considered enriched.

Whole-body metabolic assessment by indirect calorimetry revealed no significant differences in respiratory exchange ratio (RER) between groups (Figure 2D–E), indicating unchanged substrate utilization. In contrast, energy expenditure (EE) was significantly increased *in S. boulardii*–treated mice compared with controls (Figure 2F–G). Consistent with these alterations in energy balance, *S. boulardii* administration was associated with a significant attenuation of body weight gain (Figure 2H–I).

No significant differences were observed in liver weight, total cecal weight, or colon length between groups (Figure 2J–L), indicating that *S. boulardii* treatment did not induce gross changes in gastrointestinal morphology. 24 hours following oral gavage, *S. boulardii* was detected in fecal samples at approximately 10^7^ CFU per gram.

#### S. boulardii reshapes gut microbial community structure without altering global diversity

Given the observed effects of *S. boulardii* on food intake, energy expenditure, and body weight, we next investigated whether these physiological changes were accompanied by alterations in gut microbial community composition and diversity.

Analysis of alpha diversity revealed no significant differences between PBS- and *S. boulardii*–treated mice across observed richness, Berger–Parker dominance, or Shannon diversity index (Figure 2A), indicating that overall species richness and community evenness were not affected by administration of *S. boulardii*. At the phylum level, relative abundance profiles were broadly comparable between groups (Figure 2B). Firmicutes and Bacteroidetes remained the dominant phyla in both conditions, with no marked shifts in overall bacterial phylum-level composition. As expected, the fungal phylum Ascomycota was significantly increased in *S. boulardii*–treated mice (Wilcoxon test, FDR-adjusted *p* < 0.05), reflecting the intestinal presence of the administered *Saccharomyces* strain. Consistent with these findings, beta diversity analysis showed no significant separation between treatment groups (PERMANOVA, 999 permutations, *p* = 0.777; Figure 2C), suggesting that *S. boulardii* administration did not induce large-scale restructuring of global microbial community composition in this model.

Despite the absence of global diversity changes, linear discriminant analysis effect size (LEfSe) identified specific taxa that were selectively altered following *S. boulardii* treatment (Figure 2D). Notably, *Bacteroides caecimuris* and several Muribaculaceae lineages were enriched in treated mice. As Muribaculaceae belong to the order Bacteroidales within the phylum Bacteroidetes, these findings indicate enrichment of multiple Bacteroidetes-associated taxa following *S. boulardii* administration. Detection of *Saccharomyces cerevisiae* confirmed sustained intestinal presence of the administered strain. In contrast, a Lachnospiraceae lineage (RGIG7193 sp910586125) was reduced.

### *S. boulardii* reshapes microbial metabolic pathways and cecal metabolite composition

To determine whether these taxonomic shifts were accompanied by functional remodeling of the microbial community, we next examined predicted metagenomic pathway profiles. Principal coordinate analysis demonstrated significant separation between PBS- and *S. boulardii*–treated mice (PERMANOVA, 999 permutations, *p* = 0.014; Figure 3A), indicating treatment-associated differences in microbial functional potential. LEfSe identified multiple pathways differentially enriched between groups (Figure 3B). Pathways enriched in *S. boulardii*–treated mice included NAD salvage, thiamine diphosphate biosynthesis (thiazole component), L-threonine biosynthesis, purine degradation, and fatty acid biosynthesis pathways, whereas pathways related to peptidoglycan biosynthesis, pentose phosphate pathway branches, CDP-diacylglycerol biosynthesis, and queuosine biosynthesis were reduced. Given previous reports that *S. boulardii* can modulate microbial pathways involved in tryptophan-derived indole production^22,Vaaben et al., 2026 – manuscript under review^, we examined genes encoding enzymes in this pathway. Differential abundance was observed for enzymes catalyzing the conversion of indole-pyruvate to indole-3-acetaldehyde, including EC 4.1.1.74 and EC 4.1.1.43 (Supplementary Figure S1). Intersection analysis further demonstrated distinct pathway sets between treatment groups (Figure 3C). *S. boulardii*–treated mice exhibited 86 additional pathways not detected in PBS controls, whereas only three pathways were unique to the PBS group. This asymmetric expansion suggests that *S. boulardii* administration was associated with a broader repertoire of predicted microbial metabolic functions, consistent with a selective restructuring of the microbial ecosystem and an increase in functional diversity.

**Figure 3.**
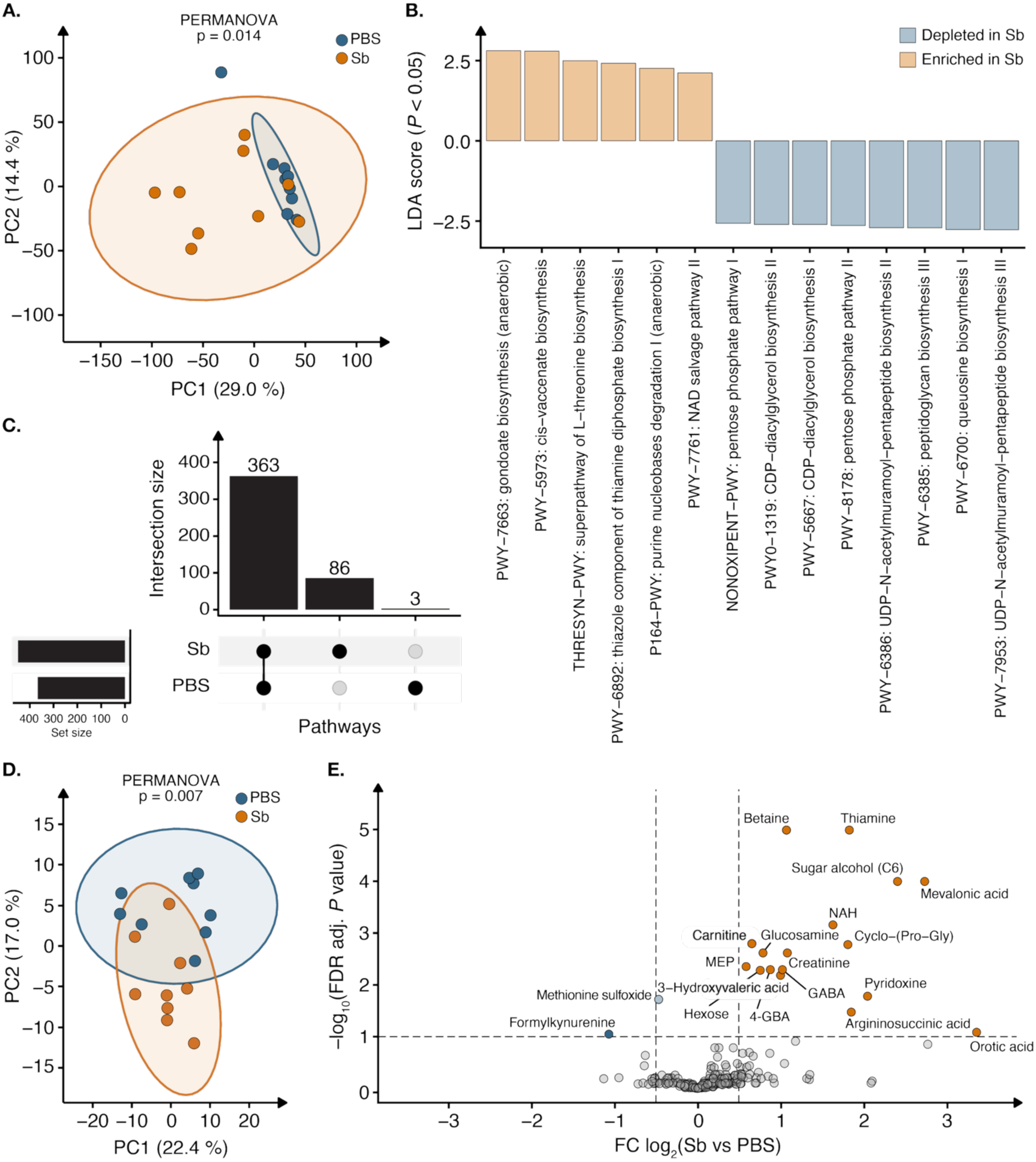
Functional metagenomic and cecal metabolomic profiling following *S. boulardii* treatment in diet-induced obese (DIO) mice. **(A)** Principal coordinate analysis (PCoA) of predicted microbial pathway profiles. Statistical significance was assessed using PERMANOVA. **(B)** Differentially enriched microbial pathways identified using linear discriminant analysis effect size (LEfSe), displayed as LDA scores indicating enrichment in PBS- or *S. boulardii*–treated mice. Group differences were tested using two-sided Kruskal–Wallis and Wilcoxon rank-sum tests with Benjamini–Hochberg false discovery rate (FDR) correction (*p* < 0.05); pathways with LDA > 2.0 were considered enriched. **(C)** UpSet plot showing pathway presence–absence relationships derived from HUMAnN3 MetaCyc annotations, indicating the number of pathways shared between or unique to treatment groups. Bar heights represent intersection sizes, and horizontal bars indicate total pathway counts per group. **(D)** Principal component analysis (PCA) of cecal metabolomic profiles. Percentage of variance explained by each principal component is indicated; statistical significance was assessed using PERMANOVA. **(E)** Volcano plot of cecal metabolites showing log_2_ fold change in *S. boulardii*–treated mice relative to PBS-treated controls versus −log_10_ FDR-adjusted *p* values. Differential abundance was defined as |log_2_ fold change| ≥ 0.5 and FDR-adjusted *p* < 0.1.

Cecal metabolomic profiling revealed a distinct metabolic signature associated with *S. boulardii* treatment. Multivariate analysis demonstrated clear separation between PBS- and *S. boulardii*–treated mice along the first two principal components, indicating a global shift in the cecal metabolome (PERMANOVA, 999 permutations, *p* = 0.007; Figure 3D). Consistent with this, differential abundance analysis identified multiple significantly altered metabolites in *S. boulardii*–treated mice (Figure 3E, Supplementary Figure S3). Among the most strongly increased compounds were betaine and thiamine, alongside additional increases in pyridoxine, GABA, glucosamine, hexose, and sugar alcohols. Several metabolites related to amino acid and nitrogen metabolism, including N-acetyl-L-histidine (NAH) and argininosuccinic acid, were also enriched following *S. boulardii* administration. In contrast, methionine sulfoxide and formylkynurenine were reduced in *S. boulardii*–treated mice, suggesting decreased oxidative or stress-associated metabolic signatures. Notably, targeted quantification of short-chain fatty acids revealed no significant differences between groups (Supplementary Figure S3).

### *S. boulardii* reshapes colonic transcriptional programs associated with inflammation and metabolism

Transcriptomic profiling of colonic tissue revealed selective, pathway-level immune and metabolic remodeling following *S. boulardii* treatment. Differential gene expression analysis comparing PBS–treated and *S. boulardii*-treated mice identified a limited number of individual transcripts reaching statistical significance (Figure 4A), indicating that *S. boulardii* did not induce broad transcriptional disruption. Notably, among the most strongly regulated genes were immune-associated transcripts, including *Igkv9-120* and *Igkv17-121*, which encode immunoglobulin κ light-chain variable regions involved in antibody diversity and B-cell–mediated immune responses, suggesting modulation of mucosal adaptive immune activity.

**Figure 4.**
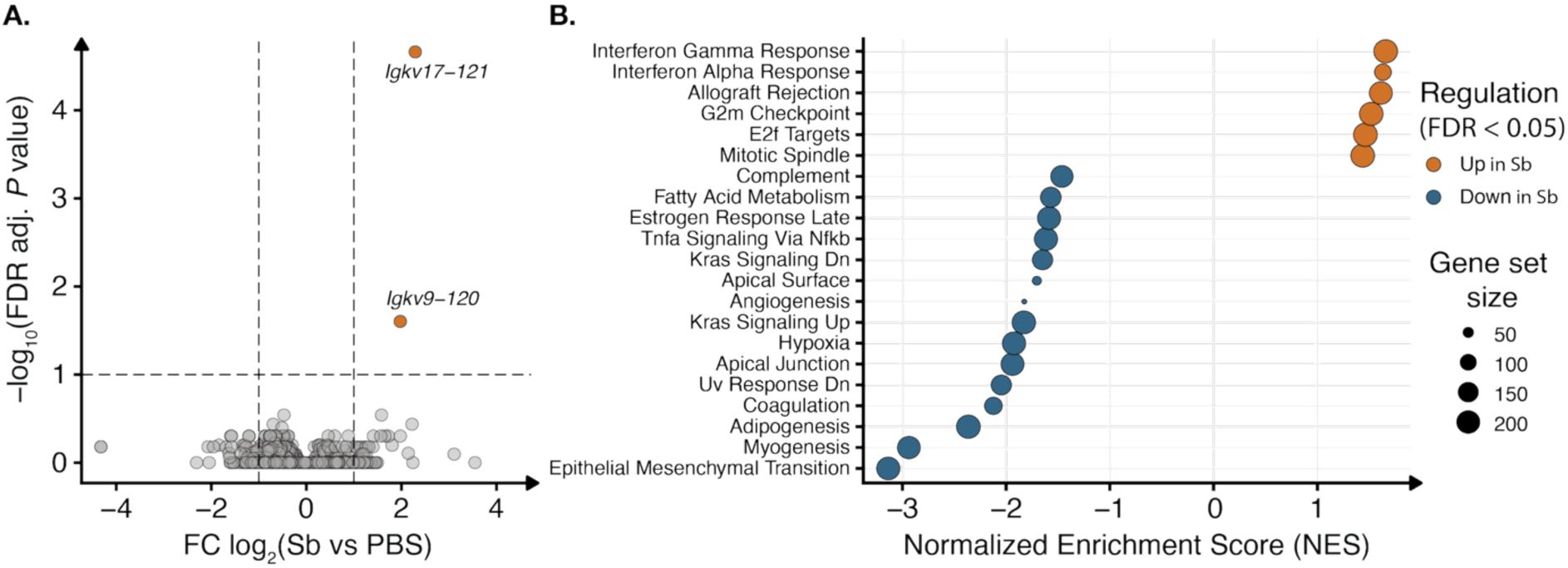
Colonic RNA-sequencing reveals pathway-level transcriptional remodeling following *S. boulardii* treatment. **(A)** Volcano plot of colonic gene expression derived from DESeq2 analysis showing *S. boulardii*–treated log_2_ fold change relative to PBS-treated mice versus −log_10_ false discovery rate (FDR)–adjusted p-values. Differential expression was defined as |log₂ fold change| ≥ 0.5 and FDR-adjusted *p* < 0.1. **(B)** Gene set enrichment analysis (GSEA) of MSigDB Hallmark pathways performed on ranked gene lists. Normalized enrichment scores (NES) are shown for pathways significantly enriched at FDR-adjusted *p* < 0.05. Dot size reflects gene set size. Positively enriched pathways indicate relative upregulation, and negatively enriched pathways indicate relative downregulation in *S. boulardi*i–treated mice compared with controls.

Although individual gene-level changes were modest, we next investigated whether *S. boulardii* treatment induced coordinated transcriptional responses at pathway level. Gene set enrichment analysis (GSEA) using the MSigDB Hallmark gene sets identified several significantly enriched biological programs (Figure 4B). This analysis revealed significant downregulation of multiple inflammation- and stress-associated pathways in *S. boulardii*–treated mice, including TNFα signaling via NF-κB, complement, coagulation, hypoxia, epithelial mesenchymal transition, and inflammatory response pathways (FDR-adjusted *p* < 0.05). In parallel, pathways related to adipogenesis and myogenesis were also negatively enriched, consistent with altered metabolic and tissue remodeling signals in the colon.

Conversely, *S. boulardii* treatment was associated with positive enrichment of interferon-α and interferon-γ response pathways, as well as pathways related to cell cycle regulation and mitotic spindle organization. These findings suggest a shift from chronic inflammatory and stress signaling toward more regulated immune surveillance and epithelial homeostasis^23^.

### *S. boulardii* modulates portal inflammatory cytokine profiles without altering circulating metabolic hormone levels

To determine whether the microbiome-, metabolomic-, and colonic transcriptomic changes induced by *S. boulardii* were reflected in systemic endocrine and immune signals, we quantified fasting portal vein hormones and cytokines. Circulating levels of key metabolic hormones, including ghrelin, GLP-1, glucagon, insulin, leptin, and PYY, were not significantly altered by *S. boulardii* treatment (Figure 5A), despite reduced food intake and attenuated body weight gain.

**Figure 5.**
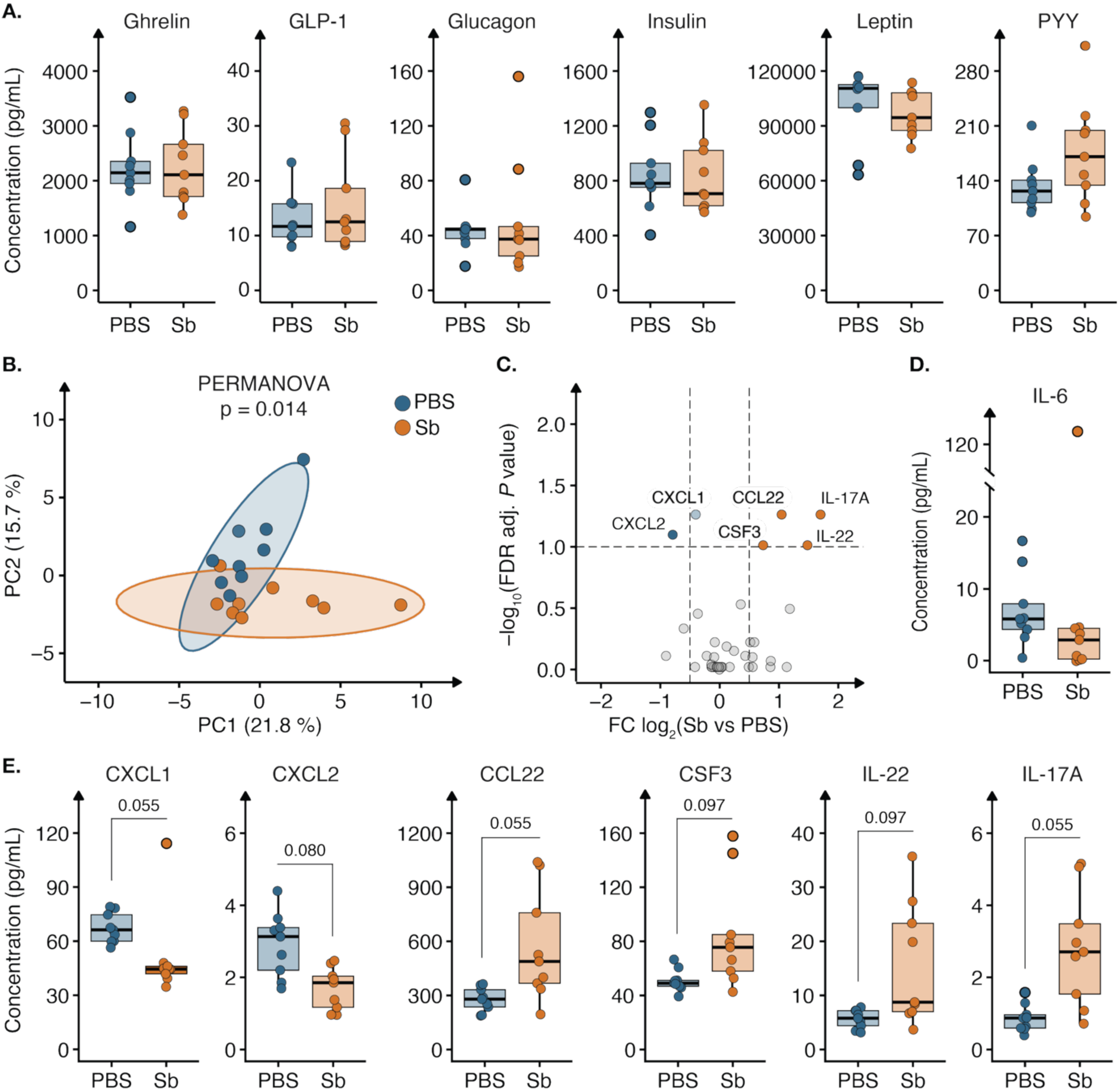
Portal cytokine and hormone profiles following *S. boulardii* treatment in diet-induced obesity mice (DIO). **(A)** Fasting portal vein concentrations (pg/mL) of metabolic hormones, including ghrelin, GLP-1, glucagon, insulin, leptin, and PYY. **(B)** Principal component analysis (PCA) of cytokine profiles. Percentage of variance explained by each principal component is indicated; statistical significance was assessed using PERMANOVA. **(C)** Volcano plot of fasting portal vein cytokines showing *S. boulardii*–treated log_2_ fold change relative to PBS-treated mice versus –log_10_ false discovery rate (FDR)–adjusted p-values. Differential abundance was defined as |log₂ fold change| ≥ 0.5 and FDR-adjusted *P* < 0.1. **(D)** Fasting portal vein concentrations (pg/mL) of IL-6. **(E)** Fasting portal vein concentrations (pg/mL) of selected cytokines, including IL-6, CXCL1, CXCL2, CCL22, CSF2, IL22 and IL-17A. Data are presented as mean ± SEM (n = 9). Statistical significance was assessed using the Wilcoxon rank-sum (Mann–Whitney U) test, with multiple testing correction applied using the Benjamini–Hochberg false discovery rate. \**P* < 0.05.

We next profiled fasting portal vein cytokines using the Olink Target 48 Mouse Cytokine panel. Principal component analysis revealed significant separation between PBS- and *S. boulardii*–treated mice (PERMANOVA, 999 permutations, *p* = 0.014; Figure 5B), indicating a treatment-associated shift in the portal inflammatory milieu. Differential abundance analysis identified several cytokines showing trends toward altered levels in *S. boulardii*–treated mice (FDR-adjusted *p* < 0.1; Figure 5C). Specifically, IL-6 exhibited a tendency toward lower concentrations, with more pronounced reductions observed for CXCL1 and CXCL2 (Figure 5D–E). In contrast, portal levels of CCL22, CSF3, IL-22, and IL-17A showed trends toward increase following *S. boulardii* administration (Figure 5E).

### *S. boulardii* is associated with coordinated metabolic–immune–microbial remodeling across multiple omics layers

To identify coordinated host–microbiome responses associated with *S. boulardii* treatment, we performed supervised multi-omics integration using Data Integration Analysis for Biomarker discovery using Latent cOmponents (DIABLO). This approach identifies sets of features across multiple omics layers that both discriminate treatment groups and covary across datasets, enabling detection of integrated molecular signatures rather than layer-specific effects.

Projection of samples onto the first two DIABLO components revealed consistent separation between PBS- and *S. boulardii*–treated mice across several data layers (Figure 6A). The clearest discrimination was observed in the metabolomics, RNA-seq, proteomics, and microbial taxa blocks, whereas physiological variables showed more moderate separation. Because DIABLO derives latent components that maximize covariance between datasets while discriminating predefined groups, these patterns indicate that multiple molecular layers contribute jointly to the treatment-associated signature.

**Figure 6.**
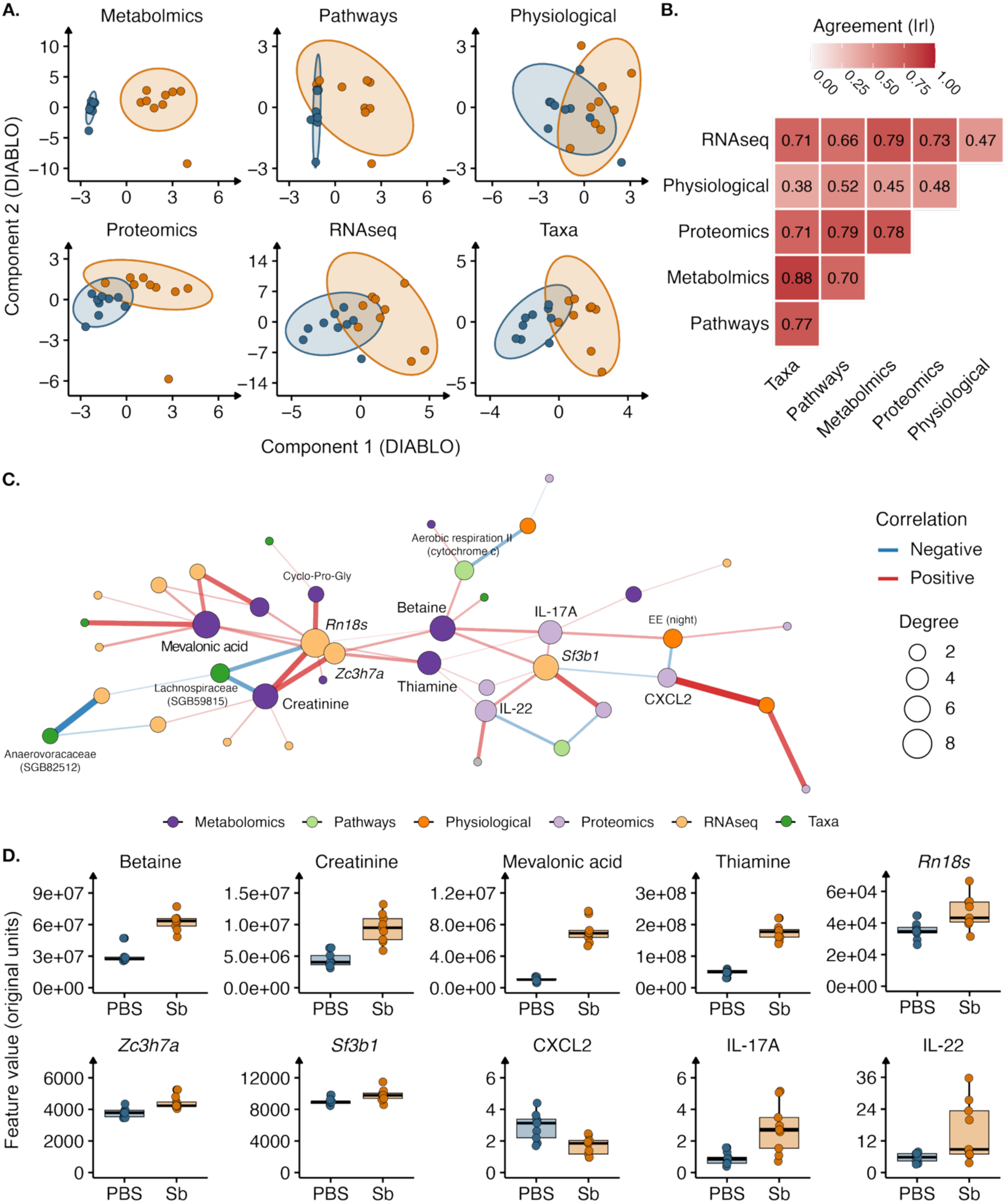
Integrated multi-omics analysis reveals coordinated cross-layer signatures associated with *S. boulardii* treatment. **(A)** DIABLO component score plots for individual data blocks (metabolomics, microbial pathways, physiological variables, proteomics (Olink), RNA-seq, and taxa) showing separation between PBS- and *S. boulardii*–treated mice along component 1 and component 2. Ellipses represent 95% confidence intervals. **(B)** Pairwise agreement (absolute correlation, |r|) between component 1 scores across data blocks. Color intensity reflects correlation strength. **(C)** Network representation of DIABLO-selected features (|ρ| ≥ 0.65) across omics layers. Nodes represent selected features colored by data type (metabolomics, pathways, physiological, proteomics, RNA-seq, taxa). Edge color indicates direction of correlation (red, positive; blue, negative), and edge thickness reflects correlation strength. Node size corresponds to degree (number of connections). **(D)** Boxplots of representative DIABLO-selected hub features across treatment groups. Data are shown as individual points with median and interquartile range.

Cross-block correlation analysis of component 1 scores demonstrated strong agreement among metabolomics, proteomics, RNA-seq, and microbial pathway blocks (|r| ≈ 0.66–0.88), indicating coordinated treatment-associated responses across molecular layers (Figure 6B). Physiological measures exhibited weaker, yet directionally consistent, correlations with these molecular blocks, suggesting that physiological changes reflect downstream manifestations of broader molecular remodeling.

Network analysis of DIABLO-selected features (|ρ| ≥ 0.65) identified an interconnected cross-omics module linking metabolites (mevalonic acid, betaine, creatinine, thiamine), inflammatory proteins (IL-17A, IL-22, CXCL2), microbial taxa (e.g., Lachnospiraceae SGB59815, Anaerovoracaceae SGB82512), and host transcripts (*Rn18s*, *Zc3h7a*, *Sf3b1*) (Figure 6C). Centrality-based hub ranking highlighted mevalonic acid, betaine, creatinine, and select RNA features as key integrative nodes bridging multiple omics layers.

Boxplot visualization of top-ranked hub features confirmed consistent treatment-associated shifts across layers (Figure 6D). *S. boulardii* treatment was associated with increased levels of mevalonic acid, betaine, thiamine, IL-17A, IL-22, and selected transcripts, alongside coordinated alterations in microbial taxa and reduced CXCL2 levels.

## Discussion

*S. boulardii* is a widely used probiotic yeast for gastrointestinal indications, including prevention of antibiotic-associated diarrhea and as adjunctive therapy for *Clostridioides difficile* infection^19,24,25^. Beyond these established uses, a recent study has shown that *S. boulardii* can alleviate cancer in colitis model through coordinated alterations in metabolic and immune pathways^26,28^, highlighting its capacity to influence host physiology beyond the gut. Furthermore, accumulating evidence indicates that *S. boulardii* exerts systemic effects on host metabolism and immune function^17,24,27,29^.

Emerging data also support a role for *S. boulardii* in metabolic disease. Experimental studies in diabetes models report improvements in glycemic control, cardiovascular and hepatic function, reductions in oxidative stress and inflammation, and favorable modulation of the intestinal microbiota^29^. Building on these observations, we demonstrate that *S. boulardii* reduces food intake, attenuates body weight gain, and increases energy expenditure in a DIO murine model. Using an integrated multi-omics approach, our study further reveals coordinated changes in gut microbial and metabolic composition, colonic transcriptional programs, and systemic immune signaling, providing mechanistic insight into how *S. boulardii* may ameliorate obesity-associated metabolic dysfunction.

Although global microbial diversity and overall community structure were largely preserved, *S. boulardii* induced selective species-level shifts accompanied by functional remodeling of the microbiome. Notably, enrichment of specific Bacteroidales lineages was observed, consistent with earlier reports describing increased abundance of Bacteroidales taxa following *S. _boulardii17,30_*_,Vaaben et al., 2026 – manuscript under review. Several enriched taxa belonged to the_ Muribaculaceae family in this study, which is frequently reduced in high-fat diet–induced obesity and associated with metabolically favorable microbial configurations in murine models^31,32^. However, in the context of prolonged high-fat diet exposure, which itself exerts strong selective pressure on microbial composition, taxonomic shifts were comparatively modest to earlier *S. boulardii* administration under chow-fed conditions^17,30,Vaaben et al., 2026 – manuscript under review^, suggesting that dietary context may constrain large-scale compositional restructuring. In addition, differences in sampling site may contribute to variation across studies, as microbial composition can differ substantially along the gastrointestinal tract^33^. Notably, the present study is based on fecal samples, whereas previous studies have analyzed cecal, colonic, or fecal contents, which may contribute to differences in the observed microbiome profiles^17,30,Vaaben et al., 2026 – manuscript under review^. The enrichment of Bacteroidales taxa, together with increased pathways related to thiamine and NAD metabolism and reduced peptidoglycan biosynthesis, therefore points toward altered microbial functional output rather than wholesale community change. Microbial structural components, including peptidoglycan and lipopolysaccharide, are known activators of innate immune receptors that promote metabolic inflammation^9,34^, a process tightly linked to NF-κB–driven signaling in obesity^35^. Reduced enrichment of such pathways may thus lower exposure to pro-inflammatory microbial signals while enhancing microbial production of cofactors and bioactive metabolites capable of influencing epithelial and immune function. Gut microbes are recognized contributors to host B-vitamin pools^36^, and the parallel increase in cecal metabolites associated with redox balance and signaling, including thiamine, pyridoxine, betaine, and GABA, supports the notion that *S. boulardii* reshapes the metabolic output of the gut ecosystem. Microbially derived neurotransmitters such as GABA have been shown to influence host neuroendocrine and immune pathways^37,38^ providing a potential link between microbial metabolism and systemic energy regulation. Consistent with previous reports linking *S. boulardii* to microbial tryptophan metabolism^22,Vaaben et al., 2026 – manuscript under review^, metagenomic pathway analysis also revealed differential abundance of enzymes catalyzing the conversion of indole-pyruvate to indole-3-acetaldehyde, a key intermediate in microbial tryptophan-derived indole biosynthesis. Although indole-derived metabolites were not significantly altered in cecal metabolite measurements, these findings suggest that *S. boulardii* may influence upstream microbial capacity for indole-derived metabolites.

The portal cytokine profile suggests that *S. boulardii* modulates immune tone at the gut–liver interface in established DIO^39^. Trends toward reduced IL-6, CXCL1, and CXCL2 are consistent with attenuation of innate inflammatory signaling linked to metabolic inflammation and align with colonic transcriptomic evidence of reduced TNFα/NF-κB–associated pathways^35^. Conversely, trends toward increased IL-17A, IL-22, and CCL22 indicate that *S. boulardii* may reshape rather than suppress mucosal immunity^40,41^. This contrasts with previous reports of reduced IL-17A following *S. boulardii* treatment ^Vaaben et al., 2026 – manuscript under review^, likely reflecting differences in experimental context. Indeed, diet-induced metabolic perturbations have been associated with both microbiota-dependent reductions in intestinal Th17 cells and IL-17A signaling, as well as impaired intestinal ILC3 function, a key component of mucosal type 17 immunity^41,42^, highlighting the context- and compartment-specific nature of type 17 immune responses. In this setting, the increase observed here may reflect partial restoration of mucosal immune activity rather than a pro-inflammatory shift. Transcriptomic enrichment of interferon response and epithelial cell cycle programs, together with differential expression of immunoglobulin-related transcripts (e.g., *Igkv17-121*), further supports remodeling of mucosal immune activity. IL-22–dependent epithelial protection^43^ and IL-17A–driven G-CSF signaling^44^ provide mechanistic context for the observed trends. Although these cytokine changes did not reach strict significance after multiple testing correction, their directional concordance (FDR-adjusted *p* < 0.1) with transcriptomic and metabolomic signatures supports a model in which *S. boulardii* rebalances gut-associated immune signaling in obesity.

Importantly, the integrated multi-omics analysis suggests that *S. boulardii* induces coordinated host–microbiome remodeling rather than isolated layer-specific effects. Increasingly, obesity and metabolic inflammation are recognized as systems-level disorders involving intertwined microbial, metabolic, and immune networks^10,35^. Multi-omics integration approaches have demonstrated that cross-layer correlations can uncover mechanistic links not evident within individual datasets^45,46^. In this context, integration across microbiome, metabolomic, transcriptomic, proteomic, and physiological datasets revealed interconnected signatures linking microbial functional pathways with cecal metabolites, host transcriptional responses, and portal cytokines. Several metabolites, including betaine, creatinine, and mevalonic acid, emerged as central nodes connecting microbial features with host immune signals such as IL-17A and IL-22. These findings support a model in which microbial alterations induced by *S. boulardii* propagate through metabolic intermediates to shape host immune signaling, highlighting the importance of cross-layer interactions in metabolic inflammation.

The strong cross-block correlations among molecular layers, together with more moderate but directionally consistent associations with physiological parameters, suggest that microbial, metabolic, and mucosal immune changes occur in a coordinated manner with shifts in host energy balance. This is consistent with emerging evidence that modulation of gut-derived inflammatory tone can influence systemic metabolism independently of major changes in circulating hormones or body weight^9,47^. Notably, while prior studies of *S. boulardii* have reported metabolic or immunological effects^17,19,20^, few have resolved coordinated host–microbe interactions at multiple molecular levels in established obesity. Our findings therefore extend the current understanding by demonstrating that *S. boulardii* acts through integrated metabolic–immune networks rather than through single-pathway modulation, providing a systems-level framework for microbiome-based intervention in obesity-associated inflammation.

Future studies should establish causality between specific microbial functions and host phenotypes, including validation of barrier integrity, immune cell composition, and tissue-specific insulin and leptin sensitivity, as well as testing the durability of metabolic effects following treatment withdrawal^48^. In particular, it will be important to disentangle direct host-modulatory effects of *S. boulardii* from indirect mechanisms mediated through shifts in microbial functional potential, as the present data indicate limited taxonomic restructuring but detectable changes in microbiome metabolic pathways. Furthermore, the lack of region-specific resolution represents a limitation, as host–diet–microbe interactions vary along the gastrointestinal tract and may differentially shape local and systemic responses^33^. From a translational perspective, the therapeutic potential of *S. boulardii* may be further enhanced through next-generation microbiome interventions, including engineered strains developed as AMTs to amplify beneficial metabolic outputs and immune-regulatory functions^13,49,50^. In conclusion, our multi-omics profiling demonstrates that *S. boulardii* modulates energy balance in established DIO and is accompanied by coordinated remodeling of the gut ecosystem, colonic inflammatory and epithelial programs, and portal immune signaling. Importantly, these effects occurred without major changes in fasting circulating metabolic hormone levels, supporting the concept that targeting obesity-associated inflammation and gut barrier–immune crosstalk may complement classical weight-loss strategies centered on appetite suppression.

## Material and method

### Animal studies

All animal procedures were approved by the Danish Animal Experiment Inspectorate (license number 2020-15-0201-00405) and conducted in compliance with Danish regulations on animal welfare. Experiments adhered to ARRIVE guidelines. Male C57BL/6JRj mice (12 weeks old; Janvier Labs) were randomized by body weight and acclimated for at least one week prior to intervention. Unless otherwise specified, mice were single-housed at room temperature under a 12-hour light/dark cycle with ad libitum access to food and water. The animal studies were carried out as single-blinded trials, with body weight and food recording measurements conducted by an animal caretaker who was blinded to treatment assignment.

All the mice were maintained on an HFD (60 %kcal) for six weeks prior to arrival and continued HFD (60 %kcal; Research Diets D12451) throughout the study. Oral gavage with *S. boulardii* was initiated at 14 weeks of age (after 8 weeks on HFD). Diet was refreshed weekly.

### Preparation and administration of *S. boulardii*

Cryopreserved *S. boulardii* (ATCC MYA-796 *ura3*Δ^51^ was streaked on yeast extract peptone dextrose (YPD) agar and incubated to obtain single colonies. Three isolated colonies were used to inoculate 5 mL YPD broth and incubated overnight at 37°C. The culture was transferred to 20 mL fresh YPD and grown for 4 hours. Subsequently, 1 mL of the culture was spread on YPD agar plates and incubated for 48 hours. Cells were harvested using 1 mL of PBS supplemented with 20% glycerol, scraped from plates, and diluted to an OD_600_ of 300 (∼3 × 10^8^ CFU).

Mice received daily oral gavage of ∼3 × 10^8^ CFU *S. boulardii* suspended in 100 µL of PBS with 20% glycerol. Control animals received vehicle only. All administrations were performed via intragastric gavage. Body weight was recorded weekly.

### Euthanasia and tissue collection

Following a 4-hour fast, mice were euthanized under anesthesia using a mixture of Hypnorm (25%) and Dormicum (25%) diluted in sterile water (0.01 mL/g body weight), followed by cervical dislocation. Blood was collected via the vena cava into lithium-heparin plasma separator tubes (PST; BD Microtainer) and processed according to the manufacturer’s instructions. Colon fecal pellets were collected for downstream microbial sequencing analyses. Liver weight, total cecal weight, and colon length were recorded at necropsy. Total cecal tissue, including luminal contents, was collected for metabolomic analyses, and distal colon tissue was harvested for transcriptomic profiling.

### Fecal sample collection

Fecal pellets were collected into pre-weighed 1.5 mL tubes containing 1 mL of PBS with 50% glycerol. Tubes were weighed post-collection to determine fecal weight. All sample processing steps were conducted at 4°C. Samples were homogenized by vortexing at 2400 rpm for 20 minutes, then centrifuged at 100 × g for 30 seconds to remove debris. Serial dilutions were plated (5 μL per dilution, in duplicate) on SC agar supplemented with uracil (20 mg/L), ampicillin (100 mg/L), kanamycin (50 mg/L), chloramphenicol (30 mg/L), and 5-fluoroorotic acid (5-FOA, 1 g/L) to inhibit bacterial and non-target yeast growth.

### Cecal content extraction

The content in provided cecum tissue samples was pressed out using a sideways motion of a stainless-steel scalpel. Cecum content was extracted in water using a constant ratio of sample mass to extraction volume. Each sample was weighed, and ultrapure water with stable isotope-labeled internal standards was added at a ratio of 1:4 (wt/wt) sample to solvent. Extracts were homogenized by beating (5 min, 30 Hz). Non-soluble material was pelleted by centrifugation (16,000 g/ 4°C/ 5 min). The supernatant was transferred to a Spin-X centrifuge tube filter (22 μm, Corning Costar) (0.22 μm) and extracts passed through using centrifugation (1500 g/ 4°C/ 5 min).

### Short-chain fatty acid (SCFA) analysis on cecal content

Short-chain fatty acid analysis was carried out by Cmbio (Vedbæk, Denmark) using gas chromatography-mass spectrometry. Samples were acidified using hydrochloric acid, and deuterium-labeled internal standards were added. Analysis was performed in randomized order using a high-polarity column (Zebron ZB-FFAP, GC Cap. Column 15 m x 0.25 mm x 0.25 µm) installed in a GC (7890B, Agilent) coupled with a quadropole MS detector (5977B, Agilent). The system was controlled by ChemStation (Agilent). Peak areas were integrated using Skyline (25.1, MacCoss Lab Software), before quantification and curation using an in-house pipeline written in MATLAB (2022b, MathWorks). Matrix effects, carryover, noise levels, and precision were evaluated using corresponding quality control samples. In case of analytical overlap of SCFA peaks with matrix contaminants, peak deconvolution was performed using Skyline 25.1 (MacCoss Lab Software).

### Semipolar metabolite analysis on cecal content

Semi-polar metabolite profiling was performed by Cmbio (Vedbæk, Denmark). The analysis was carried out using a Vanquish LC (Thermo Fisher Scientific) coupled to a Orbitrap Exploris 240 MS (Thermo Fisher Scientific). The UHPLC used an adapted method described by Doneanu et al. (UPLC/MS Monitoring of Water-Soluble Vitamin Bs in Cell Culture Media in Minutes, Water Application note 2011, 720004042en). An electrospray ionization interface was used as an ionization source. Analysis was performed in positive and negative ionization mode under polarity switching.

Untargeted data processing Metabolomics processing was performed untargeted using Compound Discoverer 3.3 (Thermo Fisher Scientific) and Skyline (25.1, MacCoss Lab Software) for peak picking and feature grouping, followed by an in-house annotation and curation pipeline written in MatLab (2022b, MathWorks). Identification of compounds were performed at four levels; Level 1: identification by retention times (compared against in-house authentic standards), accurate mass (with an accepted deviation of 3ppm), and MS/MS spectra, Level 2a: identification by retention times (compared against in-house authentic standards), accurate mass (with an accepted deviation of 3ppm). Level 2b: identification by accurate mass (with an accepted deviation of 3ppm), and MS/MS spectra, Level 3: identification by accurate mass alone (with an accepted deviation of 3ppm). Annotations on level 2b are based on accurate mass and MS/MS spectra measured with high resolution Orbitrap ESI-MS in mzCloud (Thermo Fisher Scientific), MassBank of North America (UC Davis) and the European MassBank (Helmholtz Centre for Environmental Research Leipzig). The annotations on level 3 are based on searches in the following libraries: Human metabolome database (version 5.0), Yeast metabolome database, ChEBI, BioCyc, *E. coli* metabolome database.

### Shotgun metagenomics on fecal content

Shotgun metagenomic sequencing was performed by Novogene Co., Ltd. using their standard library preparation and sequencing procedures. Briefly, genomic DNA was extracted from each sample and randomly fragmented. Fragmented DNA underwent end repair and A-tailing, followed by ligation of Illumina-compatible sequencing adapters. Libraries were subsequently size-selected and purified, and their quality was evaluated using Qubit fluorometry, quantitative PCR, and fragment size analysis. Libraries meeting quality criteria were pooled based on effective concentrations and sequenced on an Illumina platform to generate 150-bp paired-end reads.

Raw sequencing reads were processed using the same quality control and host-decontamination workflow as in our established metagenomic pipeline. In brief, adapter sequences and low-quality bases were trimmed using KneadData (v0.12.1), and reads not meeting quality and length thresholds were removed. Host-derived sequences were identified by alignment to the mouse reference genome (GRCm38) using Bowtie2 and subsequently discarded. The resulting high-quality, host-depleted reads were retained for downstream taxonomic and functional analyses.

Taxonomic profiling was performed using MetaPhlAn (v4.0). Sample identifiers were harmonized with study metadata, and analyses were restricted to samples from the PBS and *S. boulardii* treatment groups. Species-level relative abundance profiles were used for diversity analyses. Alpha diversity metrics, including observed richness, Shannon entropy, and Berger–Parker dominance, were calculated in R using the vegan package. Differences between groups were assessed using Kruskal–Wallis tests with Benjamini–Hochberg false discovery rate (FDR) correction. Beta diversity was assessed using Bray–Curtis dissimilarities, visualized by principal coordinates analysis (PCoA), and tested using PERMANOVA (adonis2). Homogeneity of dispersion was evaluated using PERMDISP.

Differential taxonomic abundance was assessed using linear discriminant analysis effect size (LEfSe) implemented in the lefser R package. Terminal taxa (leaf nodes) from the MetaPhlAn taxonomic hierarchy were retained and converted to relative abundances prior to analysis. LEfSe was performed using treatment group as the class variable, with the PBS group as the reference level, applying Kruskal–Wallis and Wilcoxon significance thresholds of 0.05 and an LDA score threshold of 2.0. Taxa meeting these criteria were considered differentially abundant.

Functional profiling was carried out using HUMAnN3. Quality-controlled reads were mapped to the ChocoPhlAn pangenome database and subsequently to the UniRef90 protein database to quantify gene families and reconstruct MetaCyc metabolic pathways. Unmapped and unintegrated features were excluded, and pathway abundances were normalized to copies per million. Analyses were performed using unstratified pathway profiles. Functional beta diversity was assessed using Aitchison distances computed from centered log-ratio (CLR)-transformed pathway data following addition of a small pseudocount, visualized by PCoA, and tested using PERMANOVA. Homogeneity of dispersion was assessed using PERMDISP.

Differential abundance of MetaCyc pathways was evaluated using LEfSe with the same statistical thresholds as for taxonomic analysis (Kruskal–Wallis *p* < 0.05, Wilcoxon *p* < 0.05, LDA > 2.0).

Presence–absence patterns of MetaCyc pathways across treatment groups were summarized using an UpSet plot generated with the ComplexUpset package. A pathway was considered present within a group if it was detected in at least one sample. In addition, targeted analyses of HUMAnN gene-family outputs were performed for selected enzyme commission (EC) functions of interest.

Multi-omics integration was performed using DIABLO implemented in the mixOmics R package. Integrated data blocks included microbial taxa, microbial pathways, cecal metabolomics, circulating proteins and hormones, physiological variables, and colonic RNA-seq data. Only samples shared across all data types were included. To avoid trivial treatment-driven separation, taxa assigned to Eukaryotes were removed prior to analysis, and the measured *S. boulardii* CFU variable was excluded from the physiological data block. Taxa were restricted to terminal nodes of the taxonomic tree, and pathway features were restricted to unstratified MetaCyc pathways. Microbial taxa and pathway data were CLR-transformed following addition of a small pseudocount and filtered for low prevalence (<10% of samples) and near-zero variance. RNA-seq data were further reduced to the 1,000 most variable genes based on standard deviation.

The DIABLO model was fitted with two latent components and a moderate inter-block design matrix. The number of retained features per block was optimized using repeated cross-validation (3-fold, multiple repeats) with centroid distance. Model performance was evaluated using repeated cross-validation, and sample separation was visualized using component score plots. Agreement between data blocks was assessed by correlations between component scores.

To characterize cross-omics relationships, features selected on the first DIABLO component were extracted and pairwise Spearman correlations were calculated across data blocks. Correlations were filtered using an absolute threshold (|ρ| ≥ 0.65) to construct an undirected network, which was visualized using tidygraph and ggraph. Nodes were annotated by data type, edge color indicated correlation direction, and node size reflected connectivity. Hub features were identified using a composite score based on degree, strength, and betweenness centrality. Key features were visualized across treatment groups using boxplots of CLR-transformed abundances.

### RNA sequencing and differential expression analysis on colon tissue

Total RNA was extracted from colon tissue of mice treated with PBS or *S. boulardii*. RNA integrity was assessed using an Agilent Bioanalyzer, and libraries were prepared using the Illumina TruSeq Stranded mRNA protocol. Paired-end sequencing (2 × 150 bp) was performed on an Illumina NovaSeq 6000 platform.

Raw sequencing reads were processed using the nf-core/rnaseq pipeline (v3.11.2) executed with Nextflow (v23.10.0) on a high-performance computing cluster using Singularity containers. Adapter trimming and quality filtering were performed within the pipeline, and quality control metrics were aggregated using MultiQC.

Reads were aligned to the *Mus musculus* reference genome (GRCm39) using HISAT2^52^. Gene annotation files provided via the --gtf parameter were used to guide spliced alignment. Aligned reads were summarized to gene-level counts using featureCounts, and the resulting count matrix was imported into R (v4.3.2) for downstream analysis. Genes with fewer than 10 total counts across all samples were excluded prior to differential expression analysis.

Differential expression analysis was performed using DESeq2 (v1.40.2). Counts were modeled using a negative binomial generalized linear model, with treatment condition (PBS vs *S. boulardii*) specified as the design variable. Statistical significance was assessed using the Wald test, and p-values were adjusted for multiple testing using the Benjamini–Hochberg method. Genes with adjusted p-values (padj) < 0.1 were considered significantly differentially expressed. Variance-stabilized expression values were obtained using the DESeq2 vst() function for visualization and exploratory analyses.

Gene set enrichment analysis (GSEA) was conducted using the fgsea package. All genes were ranked without significance filtering by the DESeq2 Wald statistic. Enrichment analysis was performed using the fgsea multilevel algorithm with gene set sizes restricted to 15–500 genes.

Mouse-native Hallmark gene sets were retrieved using msigdbr (*M. musculus*) and analyzed using Ensembl gene identifiers. Pathways with adjusted *p*-values < 0.05 were considered significantly enriched, and normalized enrichment scores (NES) were used to assess the direction and magnitude of pathway regulation.

### Plasma cytokine and hormone quantification

Plasma cytokine levels were quantified using proximity extension assay (PEA) technology (Olink Proteomics) with the Olink Target 48 Mouse Cytokine panel, according to the manufacturer’s instructions. Plasma hormone concentrations were measured using electrochemi-luminescence-based multiplex assays (Meso Scale Discovery, MSD) with the U-PLEX Metabolic Group 1 (Mouse) panel, according to the manufacturer’s instructions. Plasma samples were diluted six-fold prior to MSD analysis. Analytes for which more than 20% of measurements fell outside the assay detection range were excluded from downstream analyses.

## Data availability statement

Metagenomic and RNA-sequencing data have been submitted to the NCBI Sequence Read Archive (SRA) and are currently pending approval; a reviewer access link will be provided as soon as it becomes available. All other data are included within the article, supplementary materials, or are available from the corresponding author upon reasonable request.

## Author contributions

Karl Alex Hedin: Conceptualization; Funding acquisition; Project administration; Data curation; Formal analysis; Investigation; Visualization; Methodology; Writing—original draft; Writing—review and editing. Troels Holger Vaaben: Data curation; Investigation; Visualization; Methodology; Formal analysis; Writing—review and editing. Ditte Olsen Lützhøft: Data curation; Methodology; Investigation; Writing—review and editing. Benjamin A. H. Jensen: Methodology; Writing—review and editing. Morten Otto Alexander Sommer: Funding acquisition; Project administration and supervision; Writing—review and editing.

## Acknowledgement

This work received funding from the Novo Nordisk Foundation under NNF Grant No. NNF20CC0035580, NNF24SA0100980, the NNF Challenge Programme CAMiT under Grant No. NNF17CO0028232 and the Distinguished Innovator Grant No. NNF220C0081058. The work also received funding from the Innovation Fund Denmark under Innoexplorer Case No. 4295-00053A.

The authors are grateful to Hitesh P. Gelli and DTU Bio Facility’s animal caretakers, Heidi Arps, Maja Danielsen, Kenneth Rene Worm, and Anders Johannes Moustgaard, for assistance with the *in vivo* experiments.

## Declaration of generative AI and AI-assisted technologies

During the preparation of this manuscript, the authors used ChatGPT-5 to improve readability and refine the language in certain sections. All content generated with these tools was subsequently reviewed and edited by the authors, who take full responsibility for the final published version.

