## Supplementary Information for "*Saccharomyces boulardii* attenuates obesity-associated inflammation and weight gain through coordinated gut ecosystem remodeling"

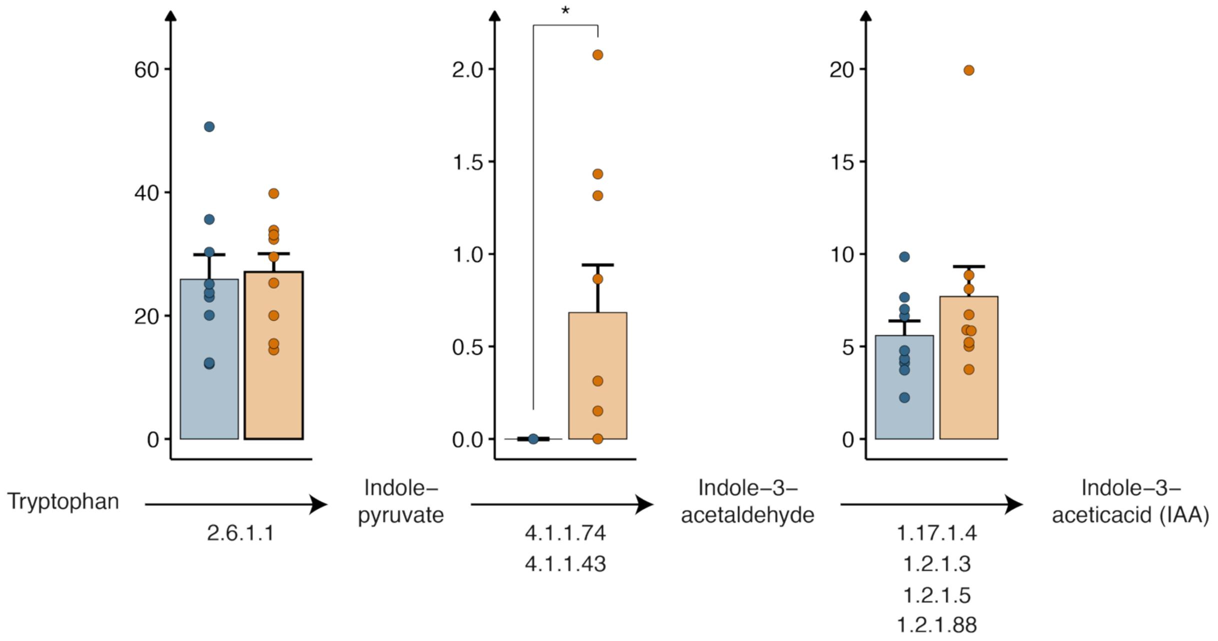


**Figure S1. Microbial pathway analysis of tryptophan-derived indole metabolism following *S. boulardii* treatment.** Bar plots showing the relative abundance of microbial genes encoding enzymes involved in tryptophan-derived indole metabolism. Enzymatic reactions are annotated with Enzyme Commission (EC) numbers and arranged to reflect the pathway from tryptophan to indole-3-acetaldehyde and indole-3-acetic acid (IAA). Bars represent mean ± SEM with individual mouse-level abundances shown as dots. Group differences were evaluated using two-sided Welch's t-tests, and multiple comparisons across EC features were corrected using the Benjamini–Hochberg false discovery rate (FDR).

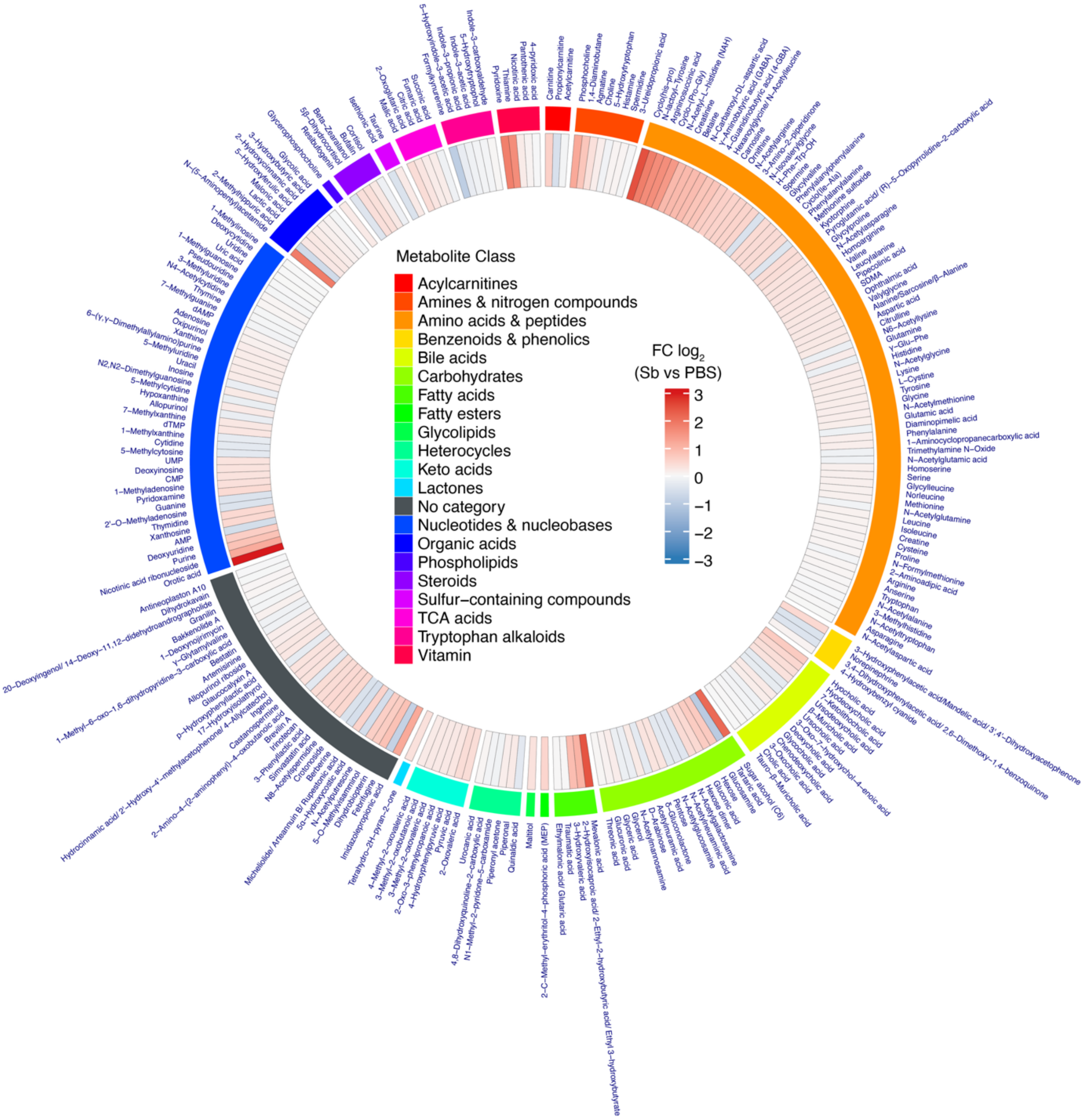


**Figure S2. Cecal metabolomic remodeling following *S. boulardii* treatment in diet-induced obese mice (DIO).** Untargeted metabolomic profiling of cecal samples from mice treated with *S. boulardii* or PBS. Metabolites are shown as $\log_2$ fold change (*S. boulardii* vs. PBS) across 248 high-confidence compounds and are color-coded by chemical class.

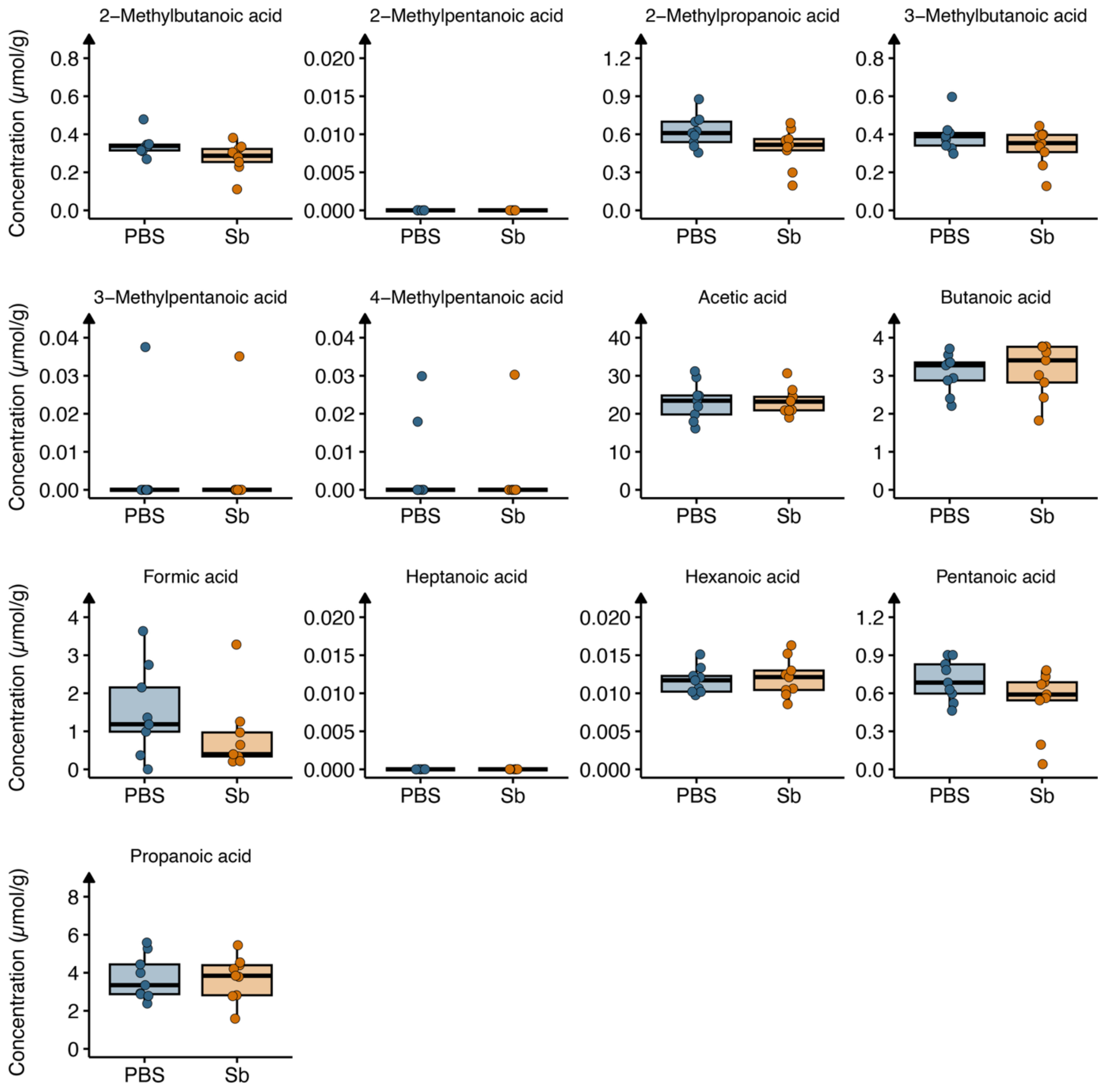


**Figure S3. Targeted quantification of cecal short-chain fatty acids following *S. boulardii* treatment in diet-induced obese (DIO) mice.** Targeted metabolomic profiling of short-chain fatty acids (SCFAs) in cecal samples from mice treated with *S. boulardii* or PBS. Concentrations are shown as absolute values (μmol/g cecal content).
